# NODeJ: an ImageJ plugin for 3D segmentation of nuclear objects

**DOI:** 10.1101/2021.11.26.470128

**Authors:** Tristan Dubos, Axel Poulet, Geoffrey Thomson, Emilie Péry, Frédéric Chausse, Christophe Tatout, Sophie Desset, Josien C. van Wolfswinkel, Yannick Jacob

## Abstract

**Background:** The three-dimensional nuclear arrangement of chromatin impacts many cellular processes operating at the DNA level in animal and plant systems. Chromatin organization is a dynamic process that can be affected by biotic and abiotic stresses. Three-dimensional imaging technology allows to follow these dynamic changes, but only a few semi-automated processing methods currently exist for quantitative analysis of the 3D chromatin organization.

**Results:** We present an automated method, Nuclear Object DetectionJ (NODeJ), developed as an imageJ plugin. This program segments and analyzes high intensity domains in nuclei from 3D images. NODeJ performs a Laplacian convolution on the mask of a nucleus to enhance the contrast of intra-nuclear objects and allows their detection. We reanalyzed public datasets and determined that NODeJ is able to accurately identify heterochromatin domains from a diverse set of *Arabidopsis thaliana* nuclei stained with DAPI or Hoechst. NODeJ is also able to detect signals in nuclei from DNA FISH experiments, allowing for the analysis of specific targets of interest.

**Conclusion and availability:** NODeJ allows for efficient automated analysis of subnuclear structures by avoiding the semi-automated steps, resulting in reduced processing time and analytical bias. NODeJ is written in Java and provided as an ImageJ plugin with a command line option to perform more high-throughput analyses. NODeJ can be downloaded from https://gitlab.com/axpoulet/image2danalysis/-/releases with source code, documentation and further information avaliable at https://gitlab.com/axpoulet/image2danalysis. The images used in this study are publicly available at https://www.brookes.ac.uk/indepth/images/ and https://doi.org/10.15454/1HSOIE.

## Background

The nucleus is a dynamic and complex structure that changes morphology and organization of its DNA content during development [1]. The spatial arrangement of chromatin within the nucleus has fundamental consequences for the accessibility and activity of regions of the genome. Changes in the location of heterochromatic domains (also known as chromocenters in *Arabidopsis thaliana* [2]) can have a drastic effect on gene expression [3] and on the maintenance of heterochromatin itself [4, 5].

Three-dimensional (3D) imaging methods are widely used to investigate nuclear morphology [6, 7]. We previously developed a workflow called NucleusJ2.0, designed to compute nuclear morphometric parameters (e.g. shape and size), as well as chromatin organization [8, 9]. While NucleusJ2.0 can automatically detect nuclei in 3D images and compute their general characteristics, further segmentation of subnuclear structures is at best a semi-automated procedure that requires user input. This induces limitations for high-throughput data analysis and for achieving consistency when processing a large number of images, while also potentially introducing user bias.

Here, we describe Nuclear Object DetectionJ (NODeJ), a new tool to automatically segment subnuclear objects such as chromocenters and Fluorescence *In Situ* Hybridization (FISH) signals. NODeJ implements an algorithm based on a Laplacian convolution [10]. This results in an increase in the contrast of objects of interest and allows to define a threshold on the enhanced image to obtain the segmented objects. The relevant parameters for the objects detected in the raw images are computed using NucleusJ2.0 methods [9] implemented within NODeJ.

## Implementation

NODeJ can be used to process images of nuclei from samples expressing fluorescent reporters, or from fixed tissues or isolated nuclei stained with DNA dyes. The program can be run as an ImageJ plugin through the graphic user interface (GUI) or the command line (CLI) mode to handle large datasets.

Our method is based on a Laplacian algorithm for object boundary detection [10]. The Laplacian method belongs to a group of mathematical methods for automated segmentation of objects based on the distribution of voxel intensities across an image. Other methods also included in this group are the watershed [11, 12] and the Isodata algorithms [13]. All of these methods use the distribution of voxel values to define the connected components of an image [10].

NODeJ assumes one nucleus per image and uses as an input the raw image of the nucleus as well as the mask of this nucleus (binary image of the nucleus) and computes the enhanced image resulting from the Laplacian operator (*δv_x_*). The voxel values of the enhanced image are used to compute a threshold value (*t*). A thresholding is then applied on the image to segment the final objects (Fig. 1).

**Figure 1.**
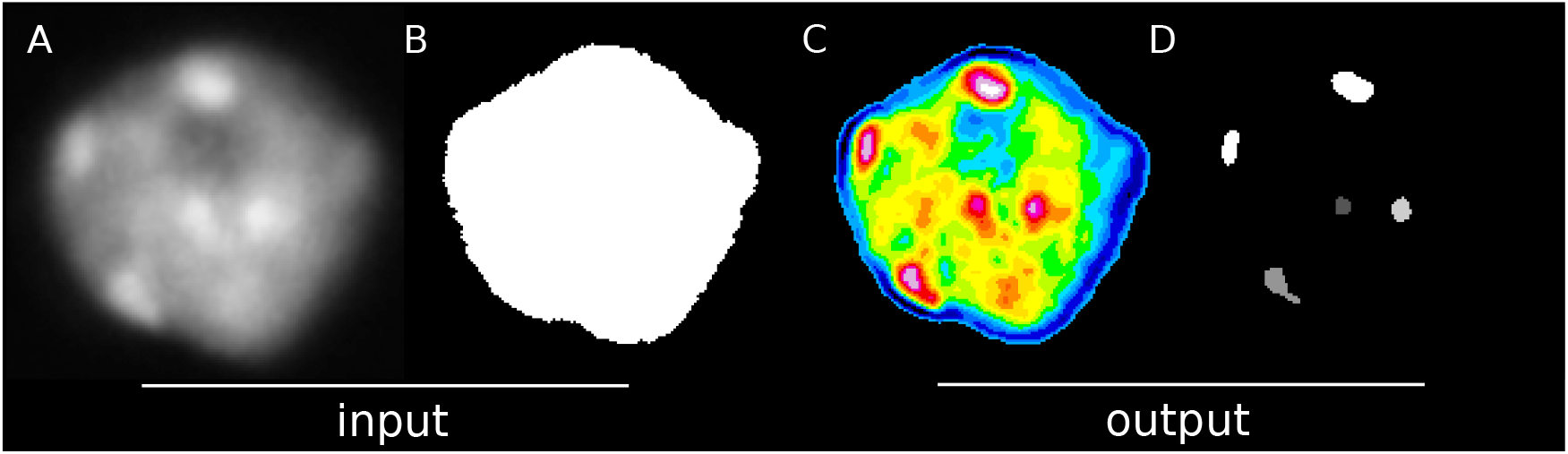
NODeJ workflow. A. Raw image of a plant nucleus (*A. thaliana*) at interphase stained with DAPI from [16]. B. Image of the segmented nucleus obtained with NucleusJ2.0. C. Image obtained with NODeJ. High voxel values are shown in red and low values are shown in blue. D. The resulting segmented image, in which each object (i.e. connected component) can be analyzed individually.

Starting from the raw image (Fig. 1A), the new voxel value is derived (Fig. 1C) for each voxel *v_x_*. NODeJ computes a locally averaged new value *δv_x_* inside the mask of the nucleus (Fig. 1B), defined as:

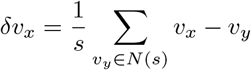

where *N*(*s*) is the neighborhood (of size *s*) of *v_x_*. The size (*s*) is defined by the user for small nuclei (volume < 50 *μm*^3^), or automatically adjusted for larger nuclei (*s* = *s* · 2.5). If the neighborhood voxel tested (*v_y_*) is outside the mask, the algorithm ignores it and goes to the next *v_y_*.

Once the enhanced image is obtained (Fig. 1C), its signal is smoothed using a Gaussian blur filter from ImageJ [14]. Then, the threshold value *t* is computed (using the enhanced image) as 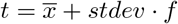, where *f* is a factor defined by the user (for the whole dataset) and automatically adjusted (*f* = *f* +1) for large nuclei (volume > 50 *μm*^3^), 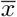 being the average of voxel values from the enhanced image and *stdev* being the standard deviation from the same image (Fig. 1C). Finally, the connected components of the binary image, obtained by applying the threshold *t*, are defined using the library morpholiBJ [15] (Fig. 1D).

## Results and discussion

NODeJ requires raw images (e.g. DNA staining [Fig. 2 and 3] or FISH [Fig. 4]) and a mask of the nuclei as input. The mask can be generated by NucleusJ2.0 (Fig. 1B). All boundaries of the nuclei need to be included in the two input images.

**Figure 2.**
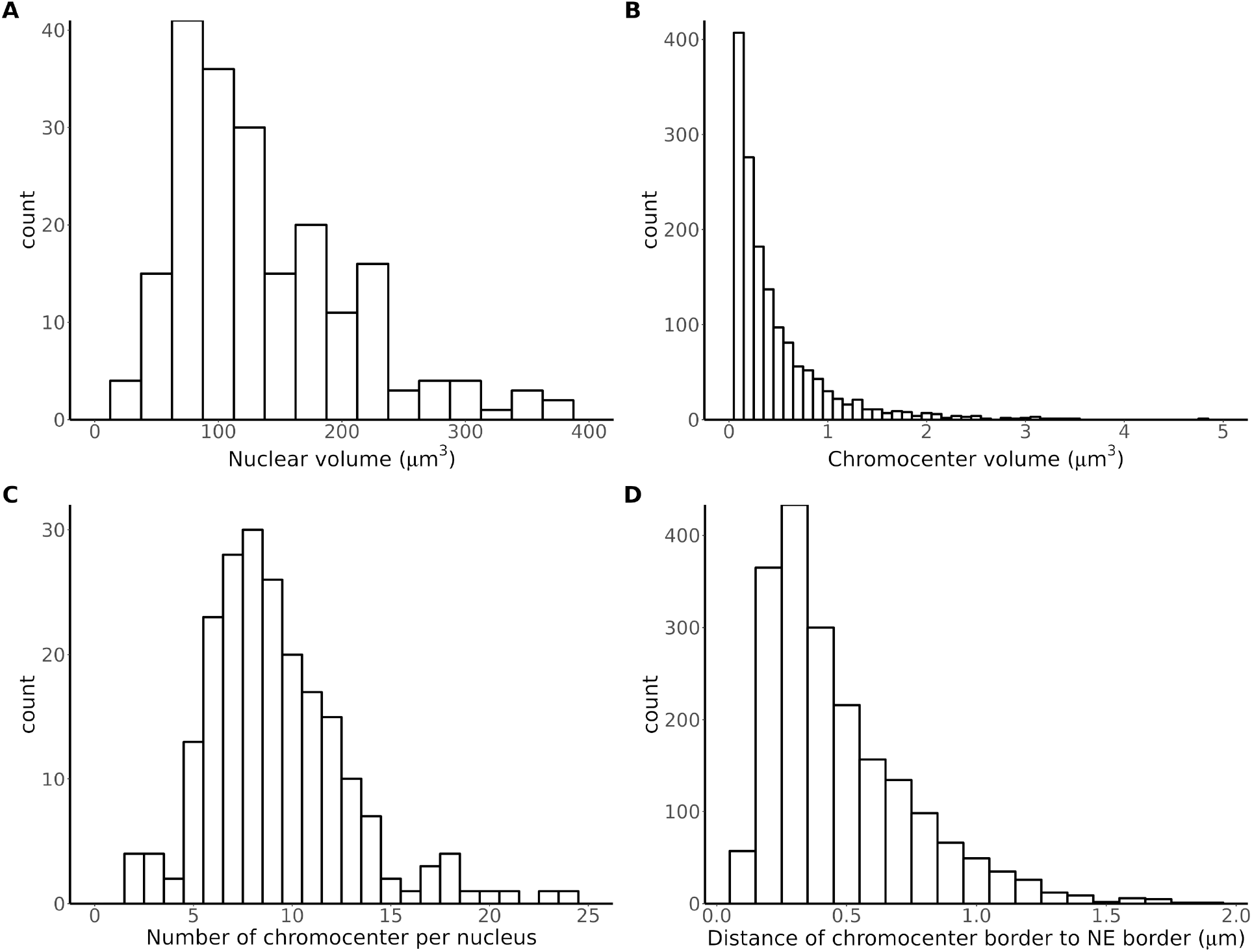
Analysis of *A. thaliana* wild type nuclei with NODeJ. Results obtained from Arpòn *et al*. (2021) describing chromocenters from isolated wild type nuclei (n=212) extracted from whole plants (Additional file 3). A to D. Histograms of chromocenters and nucleus characteristics. The histograms show the repartition of nuclear volume (A), the chromocenter volume (B), the chromocenter number by nucleus (C) and their distance to the nuclear envelope (D). The segmentation of the nuclei were obtained using NucleusJ2.0 with default parameters [9]. Histograms were made using various R packages [20, 21] (Supplementary tables 1 and 2 describe the computed parameters).

**Figure 3.**
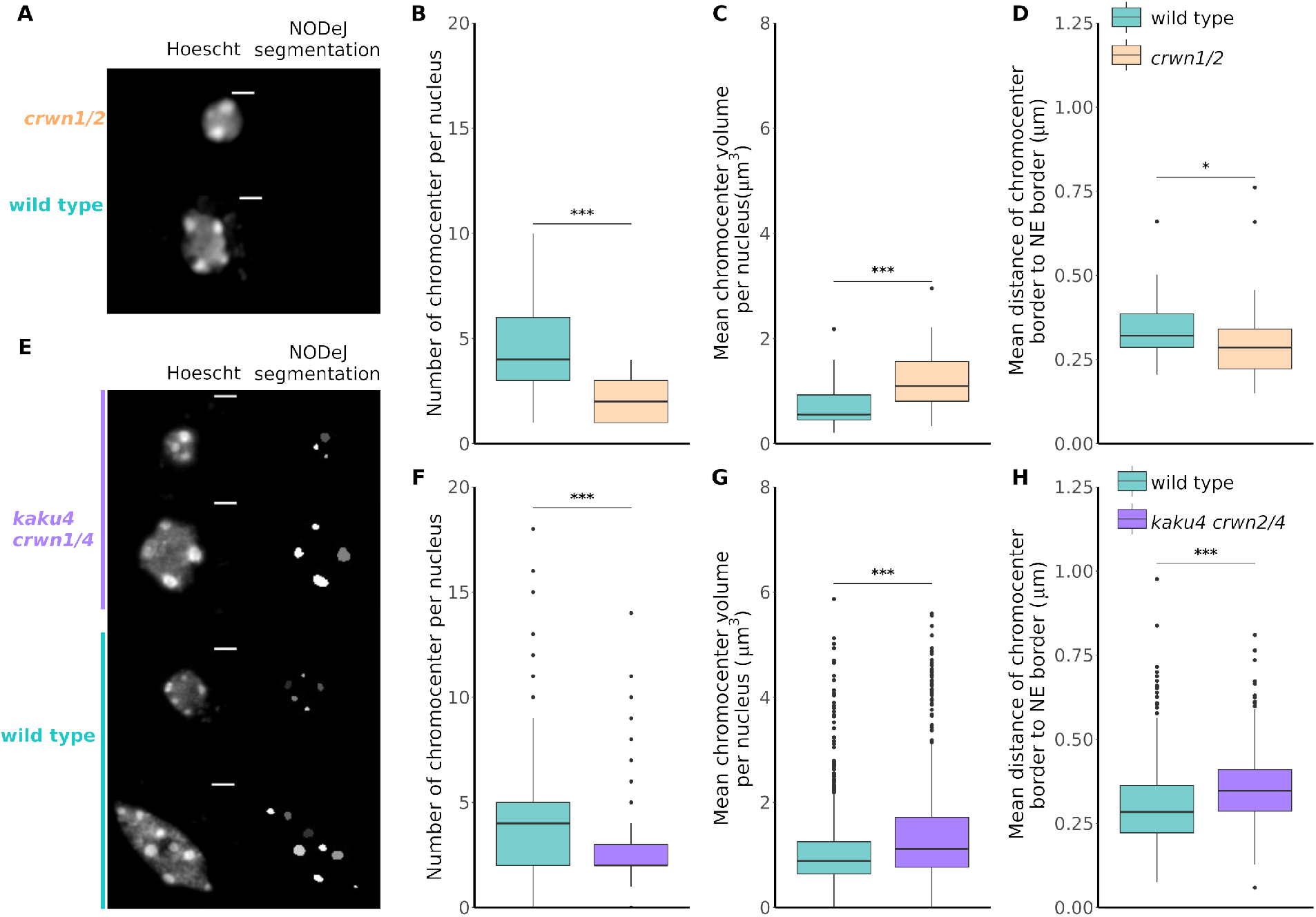
NODeJ analyses of two datasets from mutants (*crwn1/2* and *kaku4 crwn1/4*) known to alter chromatin organization in *A. thaliana* cotyledon epidermis. A. Z-projection of guard cell nuclei (diploid cell) of wild type and *crwn1/2* mutant leaf epidermis, stained with Hoechst as well as the NODeJ image result (scale bar 2 *μm*). *crwn1/2* mutants (n=39) and wild type plants (n=38) from Poulet *et al*. (2015) (Additional file 3). B. Number of chromocenters. C. Mean chromocenter volume per nucleus. D. Mean distance from chromocenter border to the nuclear envelope per nucleus. E. Z-projection of epidermis (diploid and polyploid cells) of wild type and *kaku4 crwn1/4* mutant, stained with Hoechst as well as the NODeJ image result (scale bar 2 *μm*). *kaku4 crwn1/4* triple mutant (n=851) and wild type plants (n=609) from Dubos *et al*. 2020 (Additional file 3). F. Number of chromocenters. G. Mean chromocenter volume per nucleus. H. Mean distance from chromocenter border to the nuclear envelope per nucleus. Mann-Whitney U test P-value: * ≤0.05, *** ≤0.001. Box plots and statistical tests were made using various R packages [20, 22, 21] (Supplementary tables 1 and 2 describe the computed parameters).

**Figure 4.**
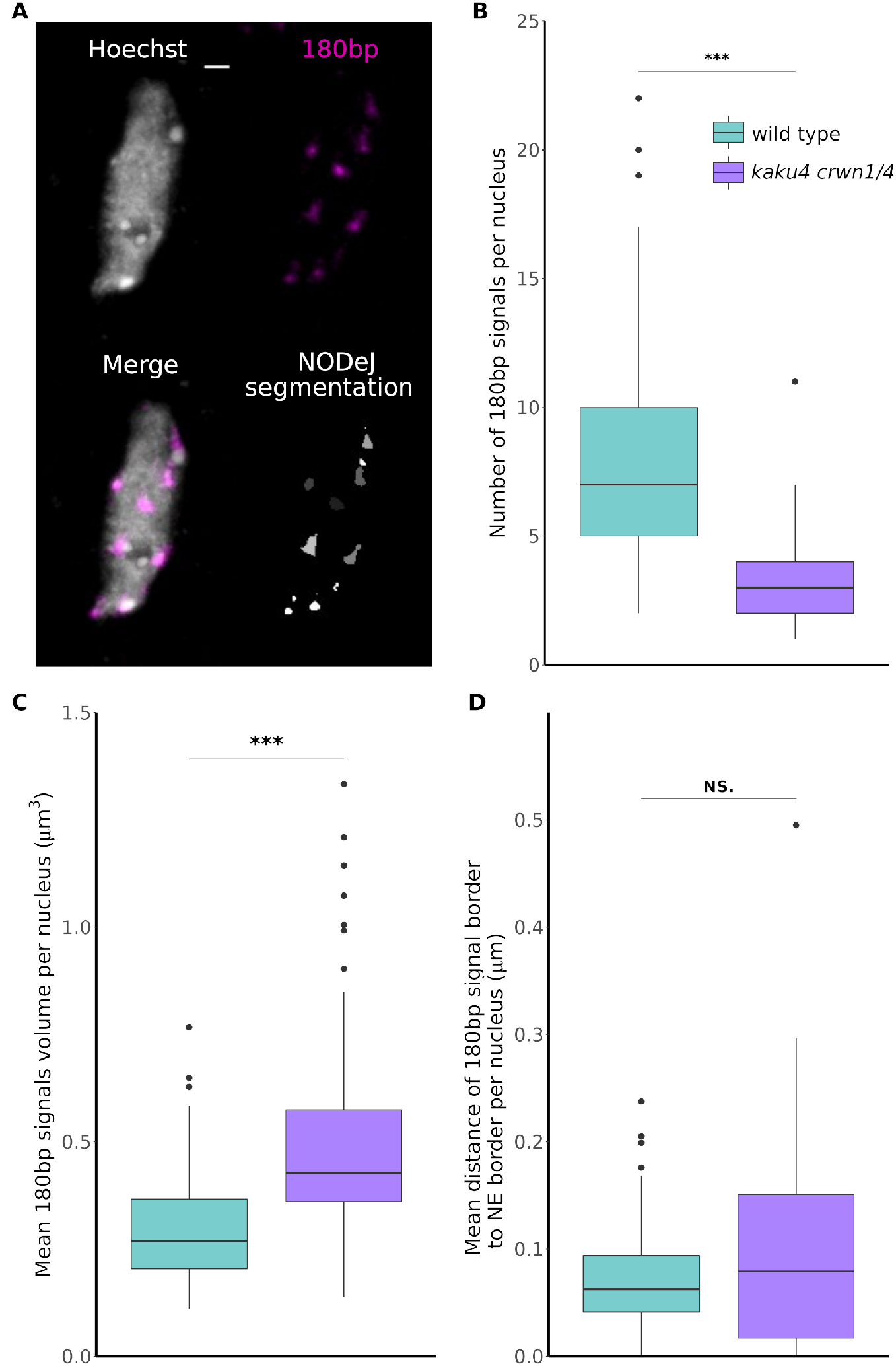
NODeJ 180bp repeat DNA FISH signal detection in cotyledon epidermis nuclei of *A. thaliana*. Results from the analysis of DNA FISH experiments of *kaku4 crwn1/4* triple mutants (n = 93) and wild type (n = 65) from the Dubos *et al*. (2020) dataset (Additional file 3). A. Z-projection of 3D DNA FISH of a wild type pavement cell nucleus (scale bar 2 *μm*). The boxplots show: B. the number of of 180bp repeat signals per nucleus, C. the mean 180bp repeat signal volume per nucleus, D. the mean distance from 180bp repeat signal border to the nuclear envelope. Mann-Whitney U test P-value: * ≤0.05, *** ≤0.001. Box plots and statistical tests were made using various R packages [20, 22, 21] (Supplementary tables 1 and 2 describe the computed parameters).

To demonstrate the performance of NODeJ, we processed two publicly-available datasets. First, we processed the images (stained with DAPI or Hoechst) from three different *A. thaliana* datasets [8, 9, 16] to identify and characterize chromocenters. Second, we analyzed centromere organization in *A. thaliana* using a DNA FISH experiment targeting the 180bp repeat, a satellite sequence from centromeres included into chromocenters [17].

### Chromocenter segmentation of 3D nuclear images from Hoechst or DAPI staining

To demonstrate the ability of NODeJ to segment objects from grayscale images, we reanalyzed several datasets of nuclei stained with DNA dyes. Arpòn *et al*. (2021) published images of isolated nuclei from various tissues from wild type plants 18 days after germination (dag) (n=212 nuclei) showing different levels of endoreduplication (i.e. ploidy level) and therefore different numbers of chromocenters. They analyzed the spacial organization of the chromocenter and showed that these heterochromatic domains are located at the nuclear periphery. This finding is in agreement with a previous report using whole-mount staining of cotyledons of *A. thaliana*, which revealed that chromocenters were localized at the nuclear periphery in the two cell types of the leaf epidermis [5].

First, we used NucleusJ2.0 [9] to obtain the masks of the nuclei from the Arpòn *et al*. (2021) dataset. We observed a similar nuclei volume distribution compared to the published results (centered ≈ 100 *μm*^3^) (Fig. 2A) [16]. We then used NODeJ to detect the size and the number of individual chromocenters for each nucleus. We found that 91% of chromocenters had a volume < 1 *μm*^3^ (Fig. 2B), and we obtained a distribution centered ≈ 8 chromocenters per nucleus (Fig. 2C), the same values found in Arpòn *et al*. (2021). We also analyzed the radial distance from the chromocenter boundary to the nearest border of the nuclear envelope. 75.5% of detected chromocenters were under 0.6 *μm* of the nuclear periphery (Fig. 2D) compared to 80% in the original study [16]. All together, the results obtained with NODeJ are in agreement with previously published results.

Next, we tested whether NODeJ is able to detect known characteristics of the nuclear periphery mutants *crwn1/2* [18] and *kaku4 crwn1/4* [9] from the datasets of Poulet *et al*. (2015) and Dubos *et al*. (2020). These mutants are known to alter nuclear morphology, chromatin organization and gene expression [18, 8, 9]. The *crwn1/2* and *kaku4 crwn1/4* mutant nuclei have a reduced number of chromocenters which are increased in volume compared to the chromocenter in wild type plants [18, 8, 9]. The nuclear images of Poulet *et al*. (2015) (39 *crwn1/2* mutant nuclei and 38 wild type nuclei) and Dubos *et al*. (2020) (851 *kaku4 crwn1/4* nuclei and 609 wild type nuclei) were acquired from the epidermis of whole-mount cotyledons (13 dag) stained with Hoechst (Fig. 3A and E). Our results show that NODeJ correctly detected the previously reported changes in heterochromatin organization in the guard cells (i.e. diploid cells forming the stomates in the leaf epidermis) of the *crwn1/2* mutant (Fig. 3B, C, D). We observed a smaller number of chromocenters (Fig. 3B), an increase of their volumes (Fig. 3C), and a slight decrease in the distance between the chromocenter periphery and the nuclear periphery (Fig. 3D) [18]. Similar results were obtained by comparing *kaku4 crwn1/4* triple mutants and wild type nuclei from the epidermis. We observed a decrease in the number of chromocenters per nucleus (average of 2.78 chromocenters per nucleus in the mutant and 4.25 for the wild type) (Fig. 3F), a 25% increase of their volumes (Fig. 3G), and an increase in the distance between the chromocenters and the nuclear envelope in the triple mutant (Fig. 3H). These differences in the *kaku4 crwn1/4* mutants obtained with NODeJ are similar to the previously published characterization of chromocenters in this mutant [9].

### 3D DNA FISH signal segmentation via NODeJ

The 180bp repeat sequence is one of the constituents of chromocenters in *A. thaliana,* and its disorganization can impact heterochromatin silencing [5]. Dubos *et al*. (2020) published a 3D image dataset of a FISH experiment labeling the 180bp repeats in *kaku4 crwn1/4* and wild type nuclei (Fig. 4A). We applied NODeJ to segment and analyze the FISH signals in these images. The results from this analysis are similar with those obtained with Hoechst staining of nuclei from the same tissue and at the same developmental stage (Fig. 3F, G, H). The number of 180bp repeat signals observed (Fig. 4B) correlated with the number of chromocenters as detected by Hoechst staining (Fig. 3F) and was reduced in the *kaku4 crwn1/4* mutant compared to the wild type. We also found an increase of the volume of the 180bp repeat FISH signals in the mutant compared to the wild type (Fig. 4C). However, we did not detect a significant change in the position of the 180bp repeats in the mutant (Fig. 4D), which differs wjem chromocenter position is assessed (Fig. 3H). Overall, the conclusions reached from using NODeJ are consitent with the results obtained with manual segmentation and analysis using NucleusJ2.0 [9].

### Comparison of object detection between NODeJ and NucleusJ2.0

The chromocenter segmentation method available in NucleusJ2.0 uses the same input files as NODeJ, but applies a watershed immersion algorithm on the raw images [11, 12]. This method in NucleusJ2.0 partitions the nucleus mask into a number of region intensities based on voxel values. The values of each region are weighed to create a map of contrast between neighbouring regions [19]. Subsequently, the user manually applies a specific thresholding to each image obtained by this method using the ImageJ GUI. This segmentation method works well to obtain a set of high intensity regions of interest to describe the nuclear chromatin organization, but the manual thresholding is slow (≈ 60 nuclei/hour). In addition, because the threshold is defined by the user, consistency between experiments, and even between nuclei, is not guaranteed. Therefore, this step represents a bottleneck for high-throughput analyses of nuclear images. In contrast, using NODeJ significantly accelerates the analysis of large nuclei datasets. For example, the analyses of the four different datasets (a total of 2014 nuclei [8, 9, 16]) using NODeJ took approximately 125 minutes of computational time, compared to ≈ 33 hours of manual labor if analyzed via NucleusJ2.0.

In order to further compare the results obtained from NODeJ and NucleusJ2.0, we used the wild type nuclei available in Poulet *et al*. (2015) and Dubos *et al*. (2020). We found that segmentation by NODeJ and NucleusJ2.0 detected similar objects, but in both datasets, the objects defined by NODeJ tended to be fewer and larger (Fig. 5). For example, we sometimes observed in the FISH analysis (Fig. 5D) that the two programs detected the same regions, but NucleusJ2.0 defined several objects whereas NODeJ only identified one object. Therefore, the number of centromeric regions expected from the DNA FISH of 180bp repeats can be overestimated in bigger nuclei by NucleusJ2.0. However, even though NucleusJ2.0 sometimes defines incorrectly subnuclear structures, this method still reveals the expected difference in the comparisons between mutants and wild type.

**Figure 5.**
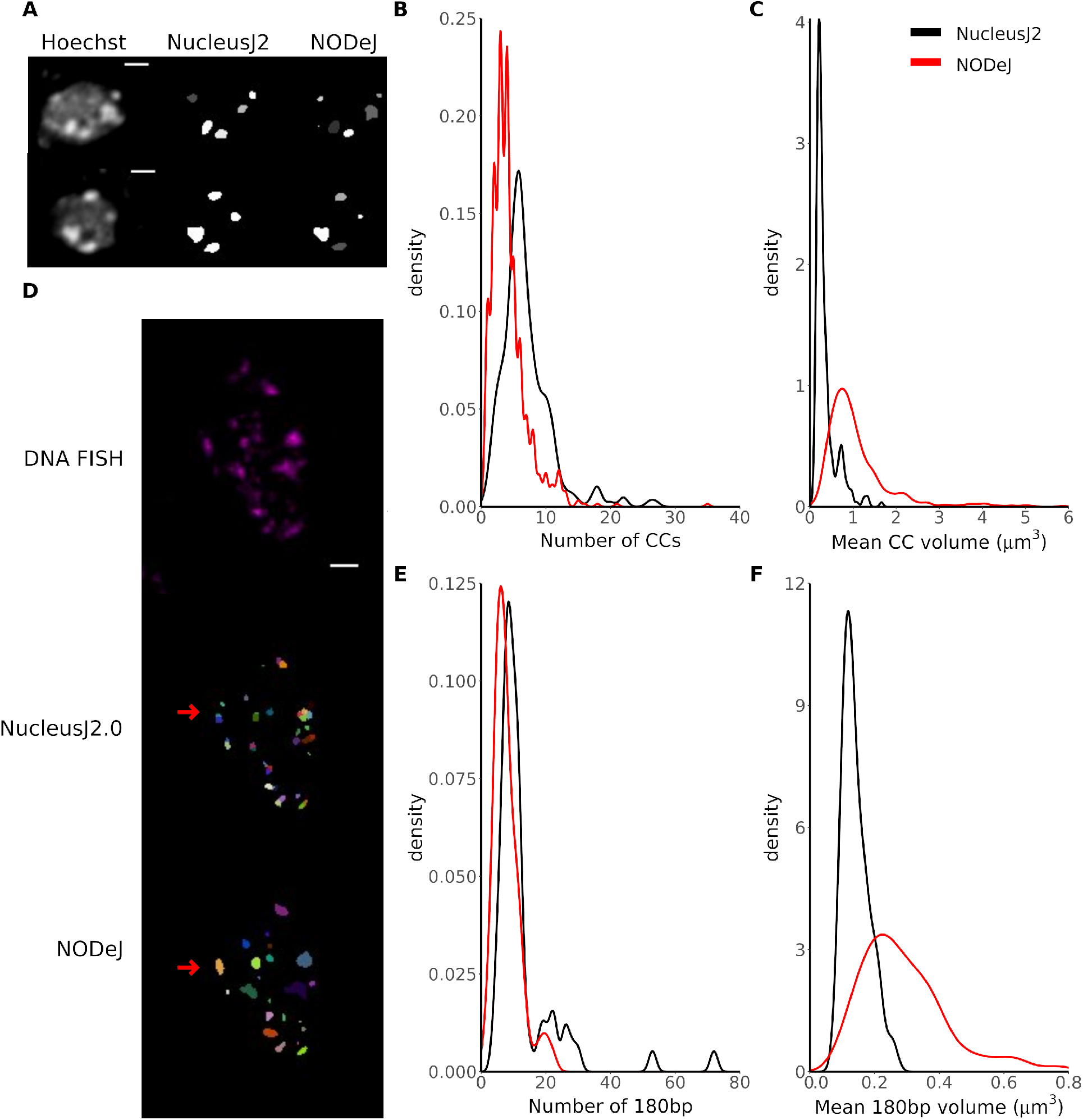
Comparison of chromocenters and FISH signals obtained with NucleusJ2.0 and NODeJ. The results were obtained from wild type nuclei from Poulet *et al*. (2015) and Dubos *et al*. (2020). We used the published results [8, 9] for the NucleusJ2.0 results (Hoechst n = 253) and reanalyzed the global set of wild type nuclei available for NODeJ results (Hoechst n = 719) (Additional file 3). A-C comparison of Hoechst staining analysis. A. Z-projection of nuclei (Hoechst staining) and NucleusJ2.0 and NODeJ segmentation result (scale bar 2 *μm*). The denstity plots show the repartition of the number of chromocenters per nucleus (B) and the mean chromocenter volume per nucleus (C). D-F comparison of FISH analysis. D. Z-projection of nuclei (180bp repeats FISH) and segmentation results from NucleusJ2.0 and NODeJ (scale bar 2 *μm*, each color is an individual object). Red arrows indicate the same region identified as two objects by NucleusJ2.0 and one object by NODeJ. The denstity plots show the repartition of the number of FISH signals per nucleus (n = 68 for NODeJ and NucleusJ2 results) (E), the mean FISH signals volume per nucleus (F). Density plots were made using various R packages [20, 21](Supplementary tables 1 and 2 describe the computed parameters).

## Conclusion

Here, we presented NODeJ, a new software tool for automated detection of subnuclear structures. We demonstrated the accuracy of NODeJ by reanalyzing three public datasets of 3D nuclear images of *A. thaliana*. NODeJ was used for detection and quantification of two types of subnuclear structures in mutants with altered chromatin organization included in these datasets. Our results are in agreement with published assessments of these *A. thaliana* mutants.

Our main conclusion is that NODeJ is an effective novel tool for the automated segmentation of nuclear images, and is able to accurately identify heterochromatic domains from a diverse set of *A. thaliana* nuclei. NODeJ was developed as an extension of NucleusJ2.0 and allows for rapid automated analysis of subnuclear structures by eliminating the semi-automated steps inherent to the use of NucleusJ2.0, resulting in reduced processing time and biases in the analysis. Our method will also be valuable for the preparation of training datasets for machine learning applications by reducing time spent during manual segmentation.

In this study, we validated the NODeJ method by analyzing characteristics of heterochromatin domains. However, we believe that the utility of this tool could be extended to other datasets such as those obtained by nucleolar labeling or even for analysis of subcellular organelles outside of the nucleus (e.g. mitochondria). The ability to extend NODeJ to additional datasets will depend on the specific subcellular structures of interest being investigated. In conclusion, our study shows that NODeJ is a valuable approach to overcome the computational bottleneck of image analysis.

## Appendix

## Acknowledgements

We thank members of our labs for discussions and advice during the course of this work.

## Abbreviations

FISH: Fluorescence *In Situ* Hybridization
3D: Three dimensional
dag: days after germination
GUI: Graphical User Interface
CLI: Command Line Interface

## Availability of data and materials

NODeJ can be downloaded from https://gitlab.com/axpoulet/image2danalysis/-/releases with source code, documentation and further information avaliable at https://gitlab.com/axpoulet/image2danalysis. The images used in this report are available in these links: https://www.brookes.ac.uk/indepth/images/ and https://doi.org/10.15454/1HSOIE.

## Author contributions

TD and AP wrote the code, ran computations, performed data analysis, and wrote the majority of the manuscript. EP, FC, CT, SD, GT, Y.J and J.C.vW made substantial contributions and revisions to the manuscript. All authors reviewed and approved the manuscript.

## Additional Files

Additional file 1: — 3D nuclear parameters

List of output parameters from NODeJ after 3D image analysis (Extracted from NucleusJ2.0 [9]) in NucAndCcParameters3D.tab file. Each line corresponds to a single nucleus.

Additional file 2 — 3D chromocenter parameters

List of output parameters from NODeJ after 3D image analysis (Extract from NucleusJ [8]) in CcParameters.tab file. Each line corresponds to one chromocenter detected.

Additional file 3 — Results of NODeJ analysis

List of output parameters obtained for each nucleus and for each chromocenter of the four datasets used in NODeJ validation [8, 9, 16]. The different parameters are explained in the Additional files 1 and 2.

