## Supplementary figures and images for "NODeJ: an ImageJ plugin for 3D segmentation of nuclear objects"

### Additional File 1

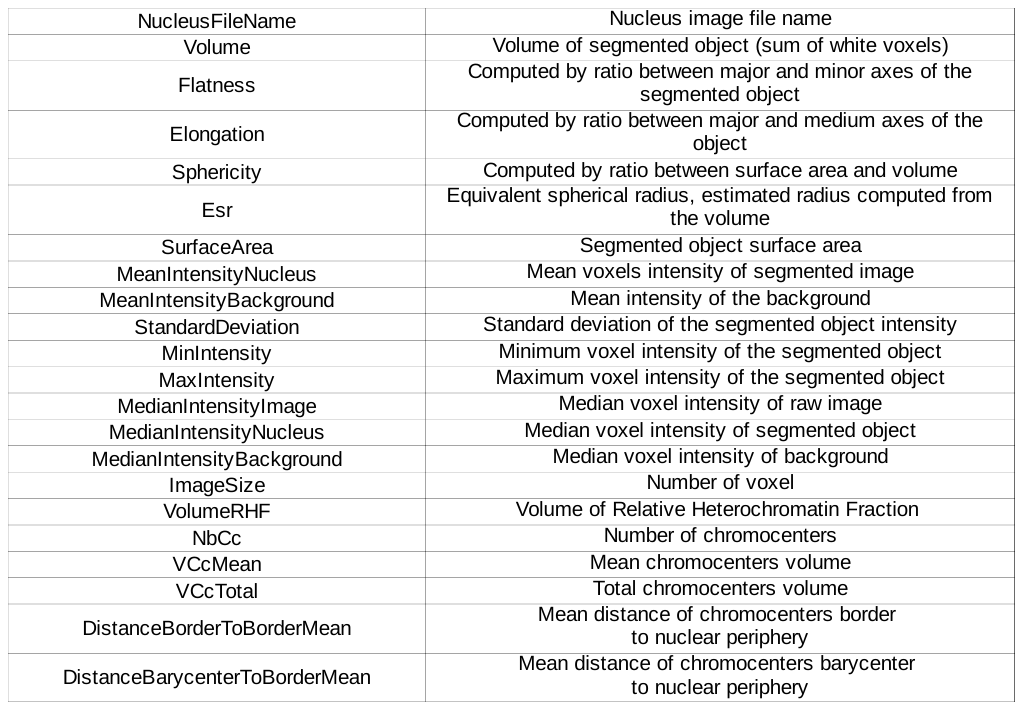

### Additional File 2

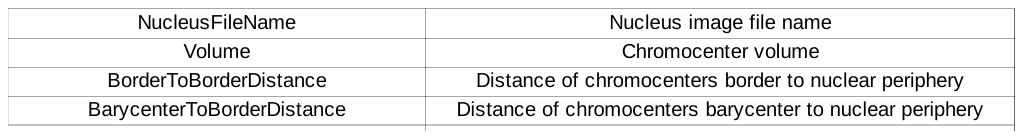
